# Network States Classification based on Local Field Potential Recordings in the Awake Mouse Neocortex

**DOI:** 10.1101/2022.02.08.479568

**Authors:** Yann Zerlaut, Stefano Zucca, Tommaso Fellin, Stefano Panzeri

## Abstract

Recent studies using intracellular recordings in awake behaving mice revealed that cortical network states, defined based on membrane potential features, modulate sensory responses and perceptual outcomes. Single cell intracellular recordings are difficult to achieve and have low yield compared to extracellular recordings of population signals, such as local field potentials (LFPs). However, it is currently unclear how to identify these behaviorally-relevant network states from the LFP. We used simultaneous LFP and intracellular recordings in the somatosensory cortex of awake mice to design and calibrate a model-based analysis method, the Network State Index (NSI), that enables network state classification from the LFP. We used the NSI to analyze the relationship between single-cell (intracellular) and population (LFP) signals over different network states of wakefulness. We found that graded levels of population signal faithfully predicted the levels of single cell depolarization in non-rhythmic regimes whereas, in delta ([2-4 Hz]) oscillatory regimes, the graded levels of rhythmicity in the LFP mapped into a stereotypical oscillatory pattern of membrane potential. Finally, we showed that the variability of network states, beyond the occurrence of slow oscillatory activity, critically shaped the average correlations between single cell and population signals. NSI-based characterization provides a ready-to-use tool to understand from LFP recordings how the modulation of local network dynamics shapes the flexibility of sensory processing during behavior.

**Significance statement:** Sensation during behaviour is strongly modulated by the animal’s internal state. Such context-dependent modulation of sensory processing is believed to largely stem from top-down control of network states in sensory cortices, with different network states being associated with distinct computational properties of the circuit. So far, a detailed characterization of network states in the awake cortex has mostly been achieved through single-cell intracellular recordings, which however cannot be easily recorded. Here, we developed a new method to classify network states from the easily accessible extracellular LFP recordings of population activity. Given the widespread use of LFPs, our work provides a critical methodology to greatly expand our understanding of the mechanisms underlying state-dependent computations in neocortex.

## Introduction

During wakefulness, behavioral and physiological markers, such as pupil diameter, whisking activity or locomotion speed vary over time and have been correlated with distinct membrane potential dynamics in rodent sensory cortices (Bennett et al., 2013; Constantinople and Bruno, 2011; Crochet and Petersen, 2006; Einstein et al., 2017; McGinley et al., 2015a; Neske et al., 2019; Nestvogel and McCormick, 2021; Okun et al., 2010; Poulet and Petersen, 2008; Poulet and Crochet, 2018; Polack et al., 2013; Reimer et al., 2014; Schiemann et al., 2015; Schneider et al., 2014). Intracellular membrane potential recordings can thus be used to provide a rich classification of how network states, defined as a set of distinctive dynamical features that include oscillations and activity or depolarization levels, change during behavior. An emergent concept for the modulation of network activity in sensory cortices is a “U-model” of network states (McGinley et al., 2015b). This model is based on the observation that, in some sensory detection tasks, performance depends on the arousal level following an inverted U-shape (it is maximal at intermediate arousal levels (McGinley et al., 2015a; Neske et al., 2019). This model posits the existence of a continuum of dynamical network states across arousal levels which includes three major and well-documented patterns of membrane potential fluctuations. At low arousal levels (low pupil diameter and absence of motor behavior), membrane potential fluctuations largely exhibit stereotypical delta-band oscillations. At moderate arousal (intermediate pupil diameter), single cells are hyperpolarized and display low amplitude membrane potential fluctuations. At high arousal levels (active motor behavior and/or high pupil dilation), the membrane potential exhibits sustained depolarization with high-frequency fluctuations and high firing activity occurs. Network states identified based on these properties of membrane potential dynamics profoundly modulate perceptual abilities and cortical processing of sensory stimuli (McGinley et al., 2015a; Neske et al., 2019).

These results shed light on how the internal state of the animal modulates sensory information processing about external stimuli. However, in many experimental settings using awake animals, extracellular measurement of population-level LFP signals is often preferred over single-cell intracellular recordings, because of low yield and high technical difficulty of intracellular experiments. Although LFPs capture subthreshold and integrative phenomena in a local neuronal population (Engel et al., 2001; Buzsáki et al., 2012; Panzeri et al., 2015), it is currently unknown how to identify the variety of network states previously described with membrane potentials from the LFP. Furthermore, while it has been reported that single-cell membrane potentials and population-level LFPs are related and their relation varies considerably across cortical states and behavioral conditions (Nestvogel and McCormick, 2021 ; Okun et al., 2010; Poulet and Petersen, 2008; Neske et al., 2019), it is not yet fully clear how to predict when they are tightly related and when they are not. Precise classification of network state variability from LFPs and its comparison with network state classification performed on membrane potentials could thus be greatly useful to enhance our understanding of how network states change during behavior and what function they may serve. Moreover, such a classification would enhance our comprehension of the relationship between single-cell and population dynamics.

By combining simultaneous intra- and extra-cellular recordings in the somatosensory cortex of awake head-fixed mice with novel analytical methods, here we developed an approach to identify, from the LFP signal alone, low-frequency rhythmic states as well as non-rhythmic network states with different levels of depolarization or hyperpolarization. We first characterized the membrane potential dynamics across different cortical states in awake mice. We then identified the LFP properties that better distinguished network states and we used those LFP properties to derive a method for robust classification of network states. Finally, we show that our classification method enables to classify well network states and explains the variability of the relationship between LFPs and membrane potentials observed across recordings of neural activity during wakefulness.

## Results

### Simultaneous intracellular and extracellular dynamics in the somatosensory cortex of awake mice: variability and network states of wakefulness

We performed simultaneous recordings of the local field potential (LFP) and the membrane potential (V_m_) of pyramidal cells in the superficial layers of the barrel cortex (S1) in head-fixed awake mice (see traces from two example recordings in Fig.1a). Recordings had a duration of 5.1±2.0 min (n=14 from 4 mice). Before proceeding to use these data to define indices of network states from the LFP, we document some of its basic properties.

**Figure 1.**
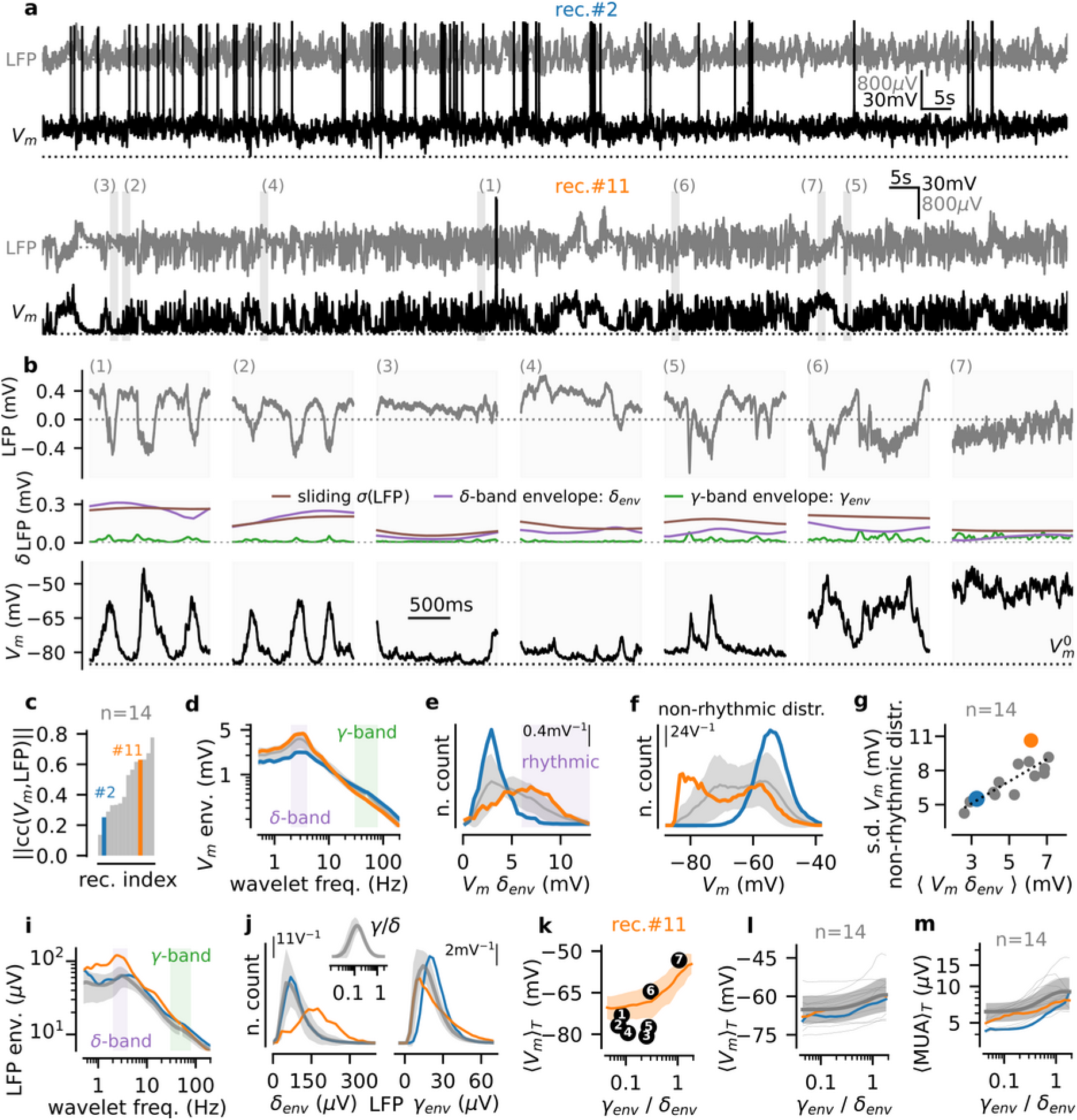
Network states of wakefulness in the mouse somatosensory cortex: electrophysiological signature and characterization based on spectral analysis. **(a)** Two example of simultaneous recordings (top: rec.#2, bottom: rec.#11) of the extracellular LFP and of the membrane potential V_m_ of a layer II/III pyramidal cell in awake mouse. **(b)** Episodes of duration 1.5 s extracted from rec.#11 at the times points highlighted in **a** (LFP on top and V_m_ on the bottom panel). In the middle panel, the time-varying standard deviation σ(LFP) evaluated over a 500 ms sliding window (brown line), the delta-envelope δ_env_ (purple line), and the gamma envelope γ_env_ (green line) of the fluctuations are shown. **(c)** Sorted histograms of the absolute correlation coefficient between LFP and V_m_ over recordings (n=14, gray). Blue (rec. #2) and orange (rec. #11) indicate the example recordings shown in **a**. **(d)** Frequency spectrum of the V_m_ signal obtained with wavelet-based time-frequency analysis (see Methods). The power-line frequency was blanked (50Hz±2Hz). We highlighted the delta (2-4 Hz) and gamma (30-80 Hz) bands in purple and green, respectively. **(e)** Histogram of V_m_ delta envelope across recordings. Time samples classified as “rhythmic” are shown in purple (see main text). **(f)** Histogram of V_m_ depolarization level for non-rhythmic samples. **(g)** Relationship between mean delta envelope of the V_m_ and standard deviation in non-rhythmic episodes per recording for all recordings. **(h)** Same as in **d** for the extracellular LFP. **(i)** Histogram of the delta envelope (δ_env_, left) and the gamma envelope (γ_env_, right) of the LFP over recordings. In the top inset, we show the histogram of the resulting gamma-to-delta ratio. **(j)** Mean depolarization level (shown as mean ± s.e.m over time samples at a given gamma-to-delta level) as a function of the gamma-to-delta ratio for a single recording (rec.#11, shown in **a** and **b**). We highlight how the gamma-to-delta measure classifies the episodes shown in **b** (see main text). **(k)** Mean depolarization level as a function of the gamma-to-delta ratio for rec.#11. **(l)** Mean depolarization level as a function of the gamma-to-delta ratio over time samples. **(m)** Relationship between Multi-Unit-Activity (MUA, see Methods) and gamma-to-delta ratio over time samples. For panels **d**,**e**,**f**, **h**, **i**, **k**, **l**, we show two example recordings (rec. #2 in blue and rec. #11 in orange) and the population data as mean ± s.e.m. over x-axis levels (n=14, gray line with shaded area).

First, we found a notable heterogeneity across recordings (compare recording #2 with #11 in Fig. 1a). In particular, the relationship between extracellular population (LFP) and intracellular (V_m_) signals was highly variable (Fig 1c, recordings were sorted by their level of absolute correlation, note the large range of observed correlation values). Second, intracellular and extracellular dynamics had a rich repertoire of activity patterns (illustrated in Fig. 1b). As previously reported under similar conditions (Poulet and Petersen, 2008; Chen et al., 2017; Einstein et al., 2017; McGinley et al., 2015a; Vinck et al., 2015), both LFP and V_m_ traces displayed epochs of rhythmic activity in the delta-band (defined as time samples with high [2,4] Hz envelope, see the V_m_ and LFP spectrums in Fig. 1d and 1i respectively, peaks were observed at 3.0±0.3 Hz for the V_m_ and 3.3±0.5 for the LFP, n=14 recordings). Those rhythmic epochs presented a high synchronization between LFP and V_m_ (correlation between V_m_ and LFP delta envelopes across time samples: 0.55±0.11, one-sample t-test for positive correlation, p=2e-10, n=14). Examples epochs of rhythmic activity are shown in traces number 1 and 2 of recording #11 (Fig. 1b). Next, confirming previous observations (McGinley et al., 2015a; Neske et al., 2019; Nestvogel and McCormick, 2021; Zerlaut et al. 2019), we found non-rhythmic epochs (here defined as time samples with a V_m_ delta envelope lower than 6mV, see Fig. 1e) at different depolarization levels (see the population histogram on Fig. 1f ranging from ~−80mV hyperpolarization levels to ~−45mV depolarization levels). Example epochs of non-rhythmic activity patterns at different depolarization levels (increasing from epoch 3 to 7) are shown in Fig. 1b. Taken together, the epochs 1-7 of Fig.1b recapitulate the different states described by the “U-model” of cortical states (McGinley et al., 2015b), that is rhythmic states of delta-band activity and non-rhythmic states at various membrane potential depolarization levels (from hyperpolarized to depolarized). Finally, recordings also differed not only in terms of the average strength of delta rhythmicity, but also in terms of the distribution of V_m_ levels over time (Fig 1f). Recordings with lower average delta envelope tended to have lower variability of V_m_ levels over time in non-rhythmic states (Fig. 1g, Pearson correlation between mean V_m_ delta envelope per recording and V_m_ standard deviation of non-rhythmic time samples, c=0.83, p=2e-4). The diversity of recordings thus filled a continuum between two qualitatively different cases of either recordings displaying mostly non-rhythmic states at an almost constant depolarization level (e.g. rec. #2 in Fig 1a, see V_m_ histogram on Fig. 1f showing a low variability of mean V_m_ over time) and recordings exhibiting overall stronger time-averaged delta V_m_ envelope and a much wider variation of the V_m_ depolarization levels (e.g. recording #11 in Fig 1a, see V_m_ histogram on Fig. 1f). These latter cases exhibited a complex dynamics with the alternation of oscillatory delta-band activity together with non-rhythmic activity at very different V_m_ depolarization levels (see example epochs of recording #11 in Fig 1b). In the next sections, we investigate how to quantitatively classify and differentiate these network states based on the extracellular LFP.

### Limitations of the existing spectral analysis of LFP for the characterization of network states with different degrees of membrane potential rhythmicity and depolarization

Because of the high impact of different strength of V_m_ rhythmicity and V_m_ depolarization levels on behavior and sensory function, we next considered how to determine a quantitative index of networks states with such V_m_ features from the more easily accessible LFP signal. Ideal properties of this index would include: i) the ability to predict the rhythmicity and depolarization of V_m_ from the LFP; ii) a U-shaped dependence of the index on depolarization levels of membrane potential and of firing of local neural populations to directly map onto the U model of network states.

We first evaluated whether existing methods based on simple spectral properties, could be used to characterize in this way, using only the LFP, the diversity of network states observed in the awake neocortex. Previous LFP-based characterization of different network states relied on the ratio between delta and gamma power (Cheng-yu et al., 2009; Saleem et al., 2010). We therefore computed the time-varying delta [2,4] Hz and gamma [30,80] Hz envelope of the LFP, and the gamma-to-delta envelope ratio over time samples (see example single recording and population histograms in Fig. 1j). We investigated the ability of the gamma-to-delta ratio to differentiate between epochs of activity that have dynamical features resembling the network states previously documented with V_m_ and described by the U-model of cortical states (McGinley et al., 2015b). To gain intuition, we first considered the example epochs 1-7, which were sorted according to the gamma-to-delta ratio of their LFP (Fig.1k, see the corresponding time-varying delta and gamma envelopes for those epochs in Fig. 1b). While this ratio could distinguish the strongly rhythmic epochs (1,2) from the high-gamma and highly depolarized non-rhythmic epoch 7 (Fig. 1k), it could not distinguish well different V_m_ depolarization levels within the non-rhythmic sets of epochs. Epoch 6 had a mean V_m_ depolarization value > 15mV higher than that of epochs 3,5, but all these 3 epochs had similar gamma-to-delta ratios (Fig. 1k). Moreover, a non-rhythmic epoch (4) had similar gamma-to-delta ratio to the two rhythmic epochs (1,2). Overall, when quantifying the dependence of mean depolarization level on the gamma-to-delta ratio across all epochs for either the example recording #11 (Fig 1k) or across all sessions (Fig 1l), it was apparent that, using the gamma-to-delta LFP ratio, it would be very difficult to distinguish between rhythmic states and non-rhythmic hyperpolarized states, and to distinguish between hyperpolarized and depolarized states within the non-rhythmic range. Next, because states with non-rhythmic and hyperpolarized V_m_ are accompanied by a reduced level of spiking activity (McGinley et al., 2015a; Neske et al., 2019; Nestvogel and McCormick, 2021; Zerlaut et al., 2019), we also analyzed the level of population firing by computing the multi-unit activity (MUA) from the extracellular recordings (see Methods). Similar to what we observed with the average V_m_ depolarization (Fig. 1l), we found that the gamma-to-delta LFP ratio had a very weak predictive power with the regard to the population spiking activity (Fig. 1m). Thus, the gamma-to-delta ratio could not be used to identify states of reduced spiking network activity during non-rhythmic activity. Reduced depolarization levels and spiking activity are important as they identify specific network states which are characterized by different properties of sensory information processing during wakefulness (McGinley et al., 2015a; Neske et al., 2019; Reimer et al., 2014; Vinck et al., 2015).

We concluded that the gamma-to-delta LFP ratio poorly differentiated rhythmic and non-rhythmic states and, more importantly, did not enable to evidence different levels of depolarization within the non-rhythmic states observed in cortical dynamics under awake condition. In the next section, we introduce a processing step of the LFP that allows such characterization.

### Using the time-varying high-gamma envelope of the LFP for a richer network state characterization

Because the membrane potential V_m_ is the reference signal for cortical state classification (Arroyo et al., 2018; Einstein et al., 2017; McGinley et al., 2015a; Polack et al., 2013; Poulet and Petersen, 2008; Reimer et al., 2014), and because we wanted to capture from the LFP the finer features of V_m_ dynamics (including the variations in rhythmicity and depolarization levels posited by the U model) that cannot be captured by the simple delta-to-gamma ratio, we next investigated whether simple mathematical transformations of the LFP displayed temporal fluctuations more qualitatively similar to those of the membrane potential.

The inverted LFP (-LFP) displays high correlation values (cc~0.5) with the membrane potential in awake animals (Poulet and Petersen, 2008; Arroyo et al., 2018) and could thus potentially provide a basis signal for the characterization of network states. However, the amplitude of the LFP is strongly dependent on the depth of the recording (Kajikawa and Schroeder, 2011; Herreras et al., 2015; Lindén et al., 2011; Sakata and Harris, 2009; Smith et al., 2012) and is subjected to drifts over short (<10s, Fig.2a, epochs (i) and (ii)) and long (>1min) time scales. These factors limit the similarity between V_m_ and - LFP and they were shown in previous work to prevent robust classification of network state during slow wave (<1Hz) activity (Mukovski et al., 2006).

**Figure 2.**
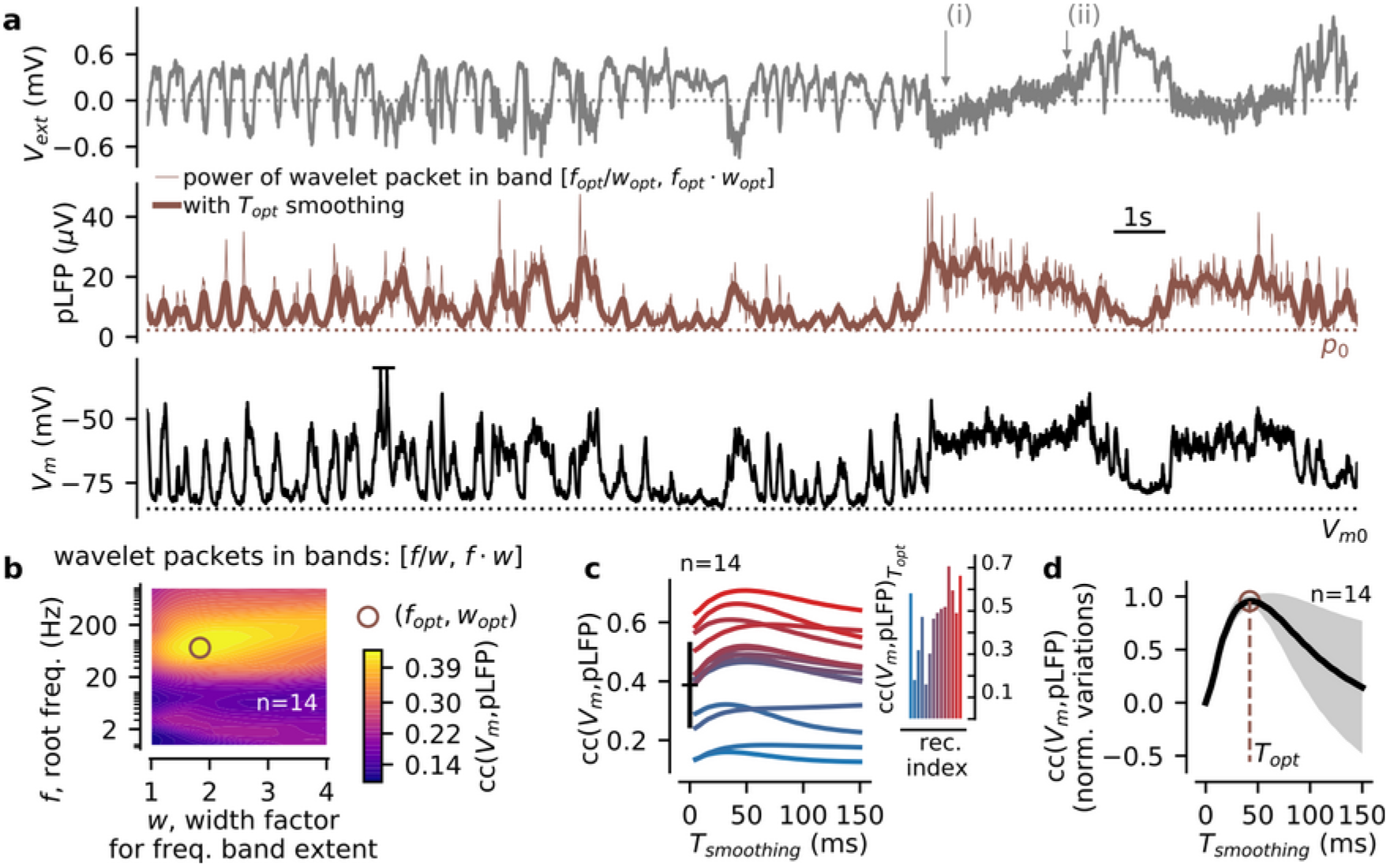
The time-varying high-gamma envelope of the LFP displays strong correlations with the membrane potential of pyramidal neurons in awake mice S1. **(a)** Example simultaneous recording of the LFP (top) and V_m_ of a layer 2/3 pyramidal cell (bottom). In the middle, we show the time-varying high-gamma envelope (brown thin line) and its smoothed fluctuations (brown thick line, the pLFP signal). The *p_0_* value (brown dotted line) corresponds to the first 100th percentile of the pLFP distribution over the whole recording. **(b)** Cross-correlation between V_m_ and the envelope of the LFP wavelet transform in the frequency band [*f/w, f·w*] (*f* is a root frequency and *w* is a width factor). We show the cross-correlation value after averaging over n=14 recordings (see the individual values per recording in **c**). Note the optimal band found for *f_opt_*=72.8Hz and *w_opt_*=1.83 (brown circle). **(c)** The effect of temporal smoothing on the cross-correlation between the LFP and *V_m_* signals. Shown for all recordings (individual recordings are color-coded according to their cc(V_m_,pLFP) value, we show the correspondence with the rec. index of Fig.1c in the inset). At *T_smoothing_*=0ms, one can see the mean value of **b** and its variability over recordings (black errorbar). **(d)** Cross-correlation between LFP and *V_m_* as function of the temporal smoothing parameter plotted after normalizing the raw cross-correlation levels of **c** by their maximum amplitude and subtraction of their level at *T_smoothing_*=0ms. With this normalization, a peak is clearly visible at *T_opt_*=42.2ms.

Guided by the previous findings in anesthetized animals (Mukovski et al., 2006), we hypothesized that the high-frequency content (*f*>40Hz, including the gamma band activity) of the LFP would provide a good predictor of the depolarization level V_m_. We therefore applied a wavelet transform to the extracellular LFP and identified the frequency band maximizing the cross-correlation with the simultaneously recorded membrane potential in our dataset (Fig.2b). This was performed by independently varying a root frequency *f* and a width factor *w*, yielding the frequency band [*f/w*, *f·w*]. For each frequency band, we divided the band into 20 evenly spaced wavelet frequencies, we computed the mean over frequencies of the wavelet envelope of the LFP (resulting in the time-varying envelope shown in Fig.2a), and we analyzed the correlation between this transformed LFP trace and the V_m_ trace after averaging over recordings (see the individual values per recording in Fig.2c sorted by recording index in the inset). We found that the band maximizing this correlation was achieved for *f_opt_*=72.8Hz and *w_opt_*=1.83, i.e. the [39.7, 133.6] Hz band (see Fig.2b). We also found that a temporal smoothing of the LFP envelope (with an optimal value *T_opt_*=42.2 ms, see Fig. 2d) enhanced its correlation with the *V_m_* signal. We refer in the following to such smoothed high-gamma envelope as the “pLFP” (processed Local Field Potential, in analogy with the terminology of Mukovski et al. (2006)). Following previous literature, we interpreted this quantity as an approximation to the time-varying recruitment of synaptic activity from a local region (diameter: ~100-200 μm) surrounding the extracellular electrode (Buzsáki et al., 2012; Gaucher et al., 2012; Katzner et al., 2009; Lindén et al., 2011; Mazzoni et al., 2011; Einevoll et al., 2013).

After this transformation of the LFP, we observed a qualitative match between the previously reported V_m_ signatures of network states (McGinley et al., 2015b) and specific features of the pLFP signal. We illustrate this finding on the recording shown in Fig.3a. We observed rhythmic activity at different envelope levels (example epochs #1,#2) as well as non-rhythmic fluctuations at various mean levels of pLFP signal (example epochs #3-#7). This similarity encouraged us to develop a quantitative network state index based on the pLFP.

**Figure 3.**
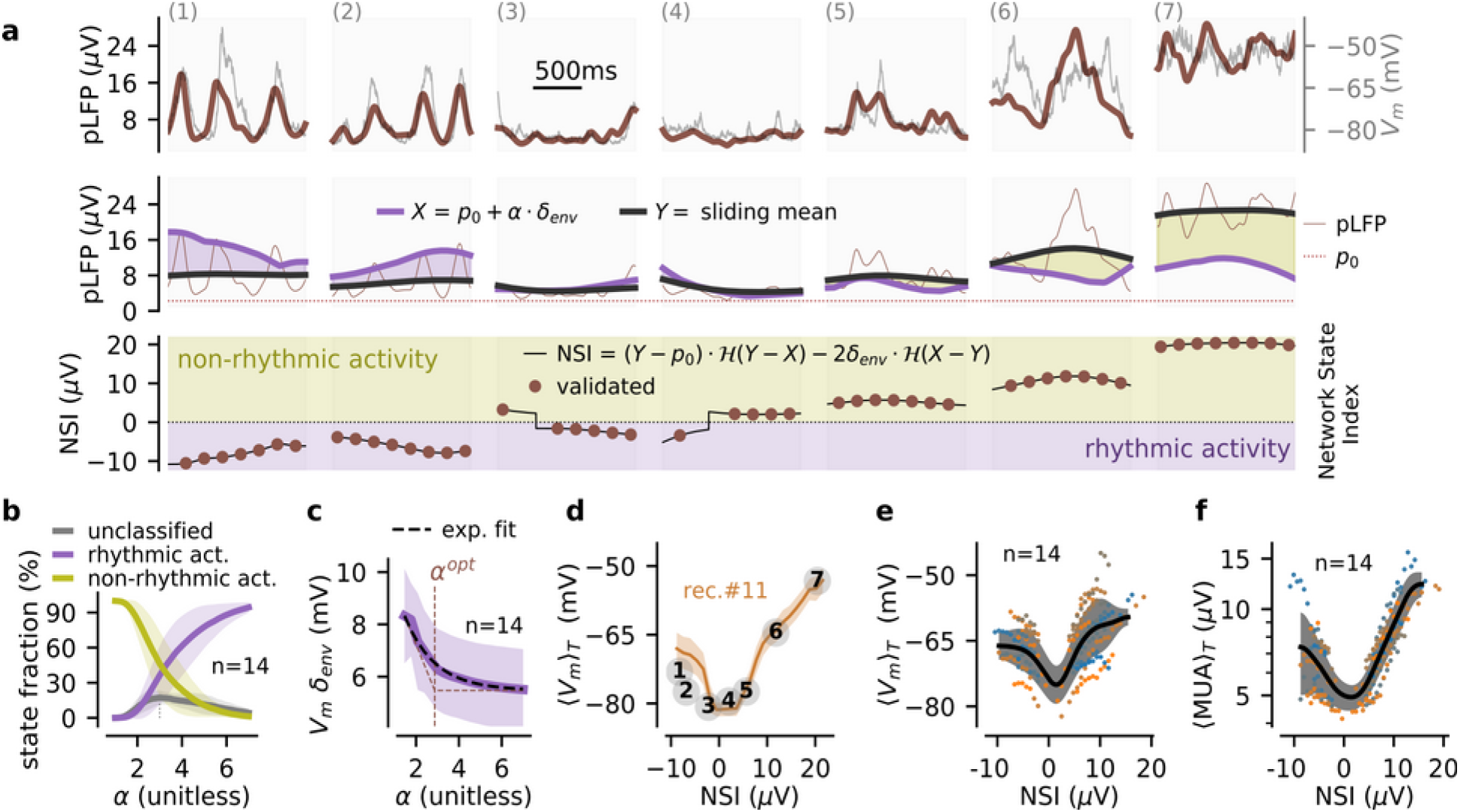
The Network State Index based on the processed LFP (NSI_pLFP_). We define a graded measure of network states based on the mean level (for non-rhythmic activity) or the low frequency envelope (for rhythmic activity) of the time-varying pLFP signal. **(a)** Example epochs of activity at different NSI_pLFP_ levels (see bottom plot), the epochs are identical to those of Fig.1b (rec. #11). In the top plot, we superimpose the pLFP fluctuations and the V_m_ fluctuations. In the middle plot, we illustrate the signal processing steps leading to the NSI_pLFP_ measure (see main text and Methods). Two time-varying quantities derived from the pLFP signal are used to classify network states: a weighted estimate of the low frequency content of the pLFP signal X(t) (purple line, shown for α=2.87) and the pLFP sliding mean Y(t) (black line). A consistency criterion validates a fraction of those as “validated” network states (brown dots). **(b)** Fraction of rhythmic, non-rhythmic and unclassified states as a function of the parameter α (weighting the propensity to classify as rhythmic states). **(c)** V_m_ envelope of the [2,4] Hz band averaged across all identified rhythmic states (mean ± s.e.m over the n=14 recordings) for different values of the parameter α. We fit the decay with an exponential function (dashed red line) and take its decay parameter as the optimal value α^opt^for the classification. **(d)** Mean depolarization level (shown as mean ± s.e.m over episodes at a given NSI_pLFP_ level) as a function of the NSI_pLFP_ measure for a single recording (rec.#11). We highlight how the NSI_pLFP_ measure classifies the episodes shown in **a** (see main text). **(e)** Mean depolarization level as a function of NSI_pLFP_ for the n=14 recordings of the dataset. We show the mean relation per recording (color-coded dots) and the mean and variability over recordings after Gaussian smoothing of 5μV width (black curve and gray area, respectively). **(f)** Relationship between Multi-Unit-Activity (MUA, see Methods) and NSI_pLFP_ over episodes for all recordings (color-code following **e**).

### Designing a Network State Index (NSI) from the processed LFP

To better discriminate specific network states of wakefulness from the LFP, we thus developed a quantitative index from the pLFP: the pLFP-based Network State Index (NSI_pLFP_). The rationale and procedure to compute the NSI from the pLFP are described in the following text and is sketched graphically with example data in Fig.3a.

To capture the slow fluctuations of network activity over time, we computed the sliding mean *Y(t)* of the pLFP over a slow time scale (*T_mean_*=500ms). Because the pLFP signal had a non-zero value at all points, we quantified the baseline of the raw pLFP signal p_0_ (set as the lowest 100^th^percentile of the pLFP distribution, and a measure of the level of baseline noise in the extracellular signal). We then analyzed pLFP fluctuations relative to this baseline level. We classified the pLFP fluctuations at such slow time scale as either rhythmic or non-rhythmic. Rhythmicity was quantified by the time-varying low frequency envelope of the pLFP fluctuations δenv(t) using a wavelet transform. We observed that establishing the rhythmic condition by only thresholding δ_env_(t) would be misleading. Indeed, the δ-band envelope was strongly co-modulated by the mean activity level Y(t) even for non-rhythmic epochs (correlation coefficients between Y(t) and δ_env_(t) in the non-rhythmic epochs defined as NSI_pLFP_>0: c=0.39±0.16 across the n=14 recordings, significance of a positive correlation: p=3.3e-10, one-sample t-test). This indicated that the δ envelope could reach high values, and cross an arbitrary threshold in absence of strong rhythmicity. This phenomenon is visible on Fig.3a: the envelope in epoch 7 is equivalent to the envelope in epoch 2 without exhibiting the clear rhythmicity in the pLFP or in the V_m_ signal that characterized epoch 2 (middle and top plots, respectively). We therefore introduced a simple model-based criterion for evaluating rhythmicity based on the following reasoning. In a noiseless, purely rhythmic setting defined by pLFP(t)=p_0_+δ_env_·(1+sin(2·π·f_δ_·t)), where δ_env_ is the envelope of the oscillation, we have Y(t)=p_0_+δ_env_. If Y(t) has an additional non-rhythmic component, we get Y(t)>p_0_+δ_env_(t) (i.e. the oscillation alone does not account for the mean level of the signal). We adapted this last relation to build our rhythmicity criterion: a higher signal mean Y(t) than the mean expected from the delta component implies non-rhythmicity. Given the non-ideal nature of the signal and to compensate for the misestimation of rhythmicity in fluctuating regimes, we rescaled the slow oscillation with a parameter α (see the next section for the determination of α), i.e. we introduced the time-varying quantity X(t)=p_0_+α·δ_env_(t). We compared the estimate of the rhythmic contribution, X(t), with the mean activity, Y(t), to quantify rhythmicity: if X(t) ≥ Y(t) activity was set as “rhythmic” (because the slow oscillation pattern is able to account for the mean activity level). Activity was defined as “non-rhythmic” otherwise. Finally, we quantified the amount of the pLFP activity in the two regimes. In the rhythmic regime, the amount of network activity was captured by the amplitude of the oscillation, 2·δ_env_(t). In the non-rhythmic regime, the pLFP deviations from baseline Y(t)-p_0_ estimated the level of ongoing activity.

The pLFP-derived NSI was defined as the amount of pLFP activity projected on the negative and positive axis for the rhythmic and non-rhythmic regimes respectively. Such a definition resulted in a continuous index where states with stronger delta components had negative values while non-rhythmic states with stronger high-gamma components had higher positive values (Fig. 3a). As highlighted by the colored area (Fig. 3a, bottom plot – purple and kaki colors for rhythmic and non-rhythmic epochs, respectively), the classification into the two states relied on the sign of the difference between the X(t) and Y(t) signals (Fig. 3a, middle plot) followed by a projection on either the negative part of the axis weighted by the oscillation amplitude 2·δ_env_(t) for rhythmic epochs, or the positive part of the axis weighted by the increase from baseline Y(t)-p_0_ non-rhythmic epochs (Fig 3a, bottom plot). The described procedure for the computation of the NSI is formalized in Eq.4 of Methods.

As the time-varying signal NSI(t) might exhibit fluctuations due to noise in both the Y(t) and X(t) quantities (X and Y are derived from the noisy LFP signal and their difference might amplify noise, see for example the signal jumps in epochs 2,3 of Fig.3a), we added a consistency criterion to NSI(t) to obtain a robust state classification of individual epochs. We first defined network state “episodes” with a window of T_state_=400ms and an update every T_state_/2=200ms. The motivation behind the choice of this time scale T_state_ was that it offered a good compromise between two constraints: T_state_ was long enough to get well defined states (e.g. more than half a cycle for oscillations in the [2,4] Hz range) and it was short enough to catch the fast and frequent switches of network states during wakefulness (McGinley et al., 2015b). The consistency criterion for episode classification required that, within a given time window of duration T_state_, the fluctuations of the NSI signal remained within a fluctuation threshold, equal to the pLFP noise level *p*_0_ (because this noise level provided an estimate of the amount of signal below which a variation is not a robust signal variation). If this stability condition was met, a network state in this time window was labelled as “validated”. The “validated” states are highlighted with brown dots over some sample epochs in Fig.3a. If this stability condition was not met, a network state at a well-defined level could not be assessed and the network state was labelled as “unclassified”. The consistency criterion prevented state validations in the presence of strong fluctuations in the time-varying NSI signal (epochs 3,4 in Fig.3a, bottom).

### Calibration of the rhythmicity threshold in the NSI definition

The parameter α in the pLFP-based NSI weights the propensity to classify the network activity as rhythmic. Its effect is illustrated in Fig.3b. Increasing α increased the proportion of rhythmic states from ~0% at α<1 to ~100% at α>6. We used simultaneous V_m_ recordings to optimize α to ensure that the classification of states as rhythmic using the LFP actually finds states that would be defined as rhythmic based on the membrane potential. In Fig.3c, we show, as a function of α, the average across all episodes classified as rhythmic of the [2,4] Hz delta envelope of the membrane potential V_m_. The delta envelope of V_m_ of the states classified as rhythmic decreased exponentially when α increased. We thus set α to a value α^opt^=2.87 that was equal to the decay constant of the V_m_ envelope as a function of α. This choice ensures we detect a large enough number of rhythmic states with a genuine amount of V_m_ rhythmicity. When classifying rhythmic states using such a value in the pLFP-based NSI algorithm, we indeed obtained that the states classified as rhythmic had V_m_ delta envelope systematically larger than the states classified as non-rhythmic (paired t-test, p=1.2e-7, n=14 recordings). Importantly, such a α value was found to be very close to the value maximizing the fraction of unclassified states (α=2.95, thin grey dashed line in Fig.3b), thus corresponding to the most conservative setting to classify network states.

### Electrophysiological signatures of NSI_pLFP_-defined network states

We then analyzed additional electrophysiological features of network regimes defined by the NSI_pLFP_ measure. We show the relationship between the NSI_pLFP_ level and the mean V_m_ depolarization value for a single recording in Fig. 3d and across all recordings in Fig. 3e. In Fig.3f, we show the relationship between the NSI_pLFP_ level and the multi-unit activity (MUA) across recordings.

States of robust rhythmicity (high δ envelope, e.g. epoch 1 in Fig.3a) showed strongly negative NSI_pLFP_ values and were associated to intermediate depolarization and MUA levels (see population data in Fig.3e,f). When the rhythmicity was not present (low values of pLFP delta-envelope, as e.g. in epochs 3,4 in Fig.3a), both the depolarization and the MUA levels had values close to their minimum (see population data in Fig.3e,f). States for which the delta component did not significantly contribute to the network activity (quantified by the pLFP) were classified as non-rhythmic (i.e. NSI_pLFP_ >0) and both their depolarization and MUA levels strongly increased with the mean level of network activity as captured by the NSI_pLFP_ (epochs 4 to 7 in Fig.3a and population data in Fig.3e,f). Importantly, the values of mean depolarization and of mean spiking activity in Fig 3e,f had a much lower standard error for any given value of the NSI_pLFP_ than the one that was found when considering the dependence of mean depolarization and of mean spiking activity on the gamma-to-delta ration (Fig 1f,1g), suggesting that the NSI_pLFP_is a much tighter predictor of membrane potential and of spiking dynamics than the gamma-to-delta ration of the LFP.

We conclude that the NSI_pLFP_, has several strengths, especially when compared to previous indices. The NSI_pLFP_ captured key features of V_m_-defined network states during wakefulness (McGinley et al., 2015b): it enabled extraction of membrane potential activity regimes ranging from delta-band activity, to asynchronous regimes at low activity levels and asynchronous regimes at high activity levels (Fig.3e,f). The NSI_pLFP_ therefore provided a quantitative measure of network states that allowed extracting the U-shape nature of cortical states from LFP recordings previously documented with intracellular recordings (Fig.3e,f).

### Evaluation of the pLFP-based NSI accuracy in estimating membrane potential based features

The above considerations suggest that it should be possible to use the NSI_pLFP_, a measure only based in LFPs, to identify reasonably well states that have either rhythmic or non-rhythmic membrane potential properties, and to identify among the states with non-rhythmic membrane potentials, those that have either depolarization or hyperpolarization of membrane potential. In this section, we quantified the accuracy of such state characterization using the pLFP-based NSI.

To this aim, we first computed the NSI on the V_m_ signal. The lower bound of the signal *p*_0_ was translated into the *V_m0_* value by taking the first percentile of the V_m_ distribution (see Fig.2a and Fig.4a). We computed the sliding mean and the time-varying low frequency envelope with the same parameters as for the pLFP signal. We derived the *X(t)* and *Y(t)* and the “V_m_-defined NSI” (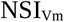) according to Eq.4 (see Methods). This V_m_- defined NSI has (by construction) negative values for the states with V_m_ rhythmicity, low positive values for states with hyperpolarized non-rhythmic V_m_, and high positive values for states with depolarized non-rhythmic V_m_. Thus, comparing the value of the V_m_-defined and pLFP-defined NSI during the validated network states enables a simple quantification of how good is the pLFP-defined NSI at identifying states of rhythmic, non-rhythmic, depolarized and hyperpolarized membrane potential.

**Figure 4.**
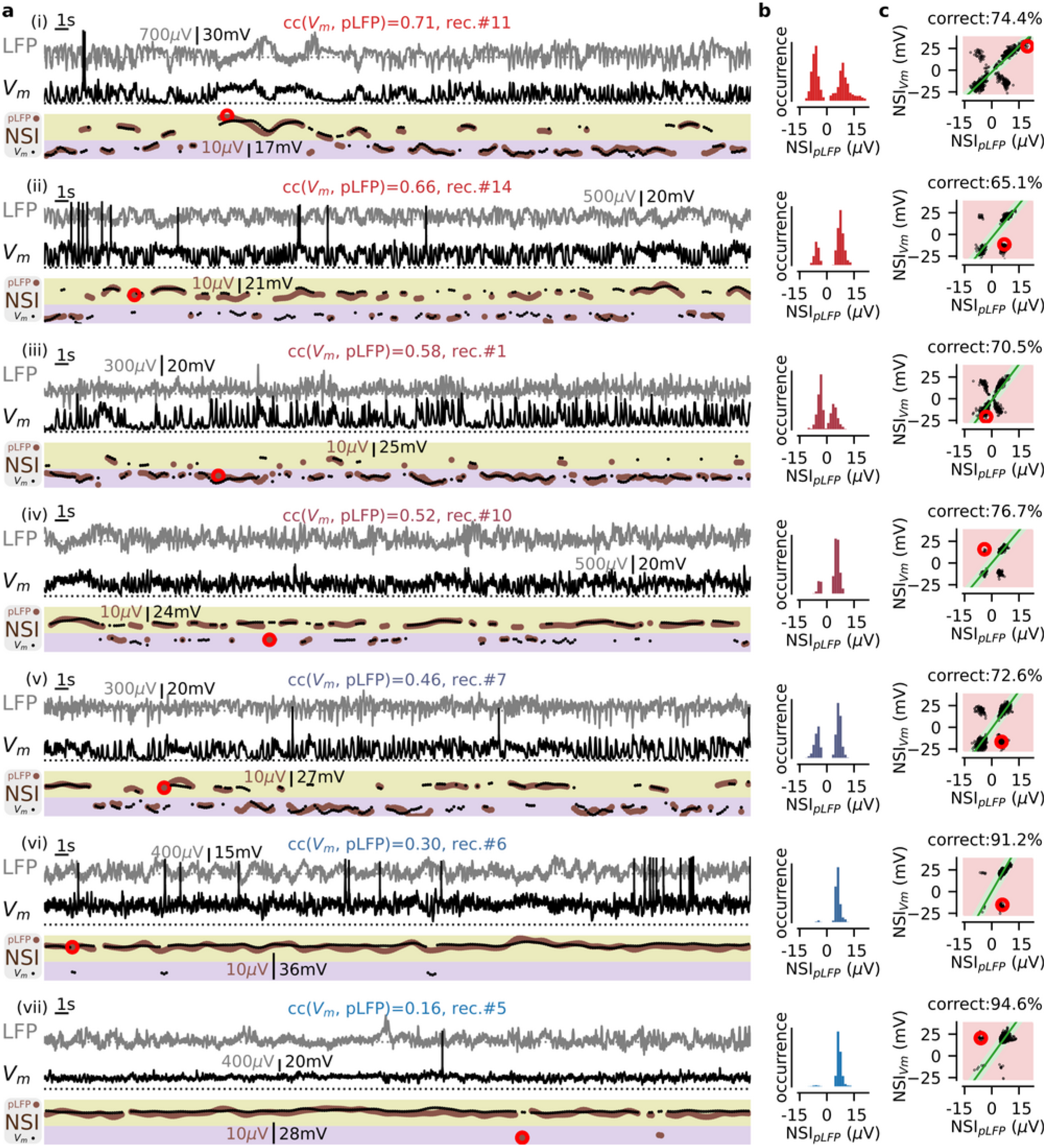
The pLFP-based NSI across multiple recordings: variability and estimated accuracy. We show 7 recordings (from i to vii) covering the whole range of observed values of correlations between the intracellular (V_m_) and extracellular (pLFP) signals. **(a)** A 60s sample of the simultaneous LFP(gray) and V_m_ (black) signals. At the bottom, we show the validated network states with their NSI value both for the “pLFP-defined NSI” (NSI_pLFP_, brown dots) and “V_m_-defined NSI” (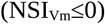, black dots). **(b)** Histogram of the NSI_pLFP_ over the whole recording length for each recording. The color code per recording (from red to blue, see also Fig.2c) represents the value of the correlation coefficient between the V_m_ and pLFP signals. **(c)** Scatter plots of the “V_m_-defined NSI” (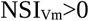) and the “pLFP-defined NSI” (NSI_pLFP_) values for all validated episodes in a given recording (i.e., extending before and after the recording sample shown in **a**). We highlight the correct area with a green color and the incorrect area with a red color. The large red circles give examples of the different sort of rejections that may happen during cross-validation (see main text).

Fig 4a shows the time course over different recording sessions of the NSI computed either on the pLFP or on V_m_. From these plots, it is apparent that the two NSI indices are remarkably well matched over the time epochs of the recordings, with the occasional presence of episodes in which the two indices were mismatched (e.g. those marked by red circles in Fig 4A). To assess when the pLFP-based NSI did and did not correctly predict the NSI measured on the membrane potential, we implemented the criterion displayed in Fig.4c. We first determined the scaling factor *F* between the V_m_-based NSI and the pLFP-based NSI, by performing the linear fit of the data predicting either both rhythmicity or both non-rhythmicity (i.e. on the lower left or the upper right of the plots in Fig.4c). This linear relationship yielded a prediction for the 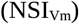 value from a NSI_pLFP_ value. Then, at a given time-point *t_i_*, the prediction was considered as correct if the difference between the predicted and observed value of 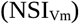 lied within a tolerance interval defined by two free parameters, *p^tol^* and 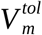. We set the value of *ptol* as the average noise level of the pLFP signal across recordings, i.e. *p_noise_*=2.85μV. For the tolerance value 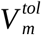, we took 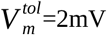 to be well above the noise level of V_m_ recordings (~0.1mV) and still have a high resolution in the [0-35] mV range of observed depolarization levels (see Fig.3e). Correct detection thus occurred when the following conditions were met: 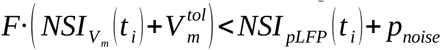 and 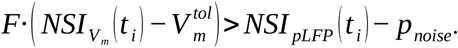. The strictness of this criterion is illustrated in Fig.4c. States were taken as incorrect when there was a mismatch between the rhythmic vs non-rhythmic classification (as shown for cases ii,iv-vii in Fig.4, see the non-matching events highlighted with a red circle). Importantly, state classification was also taken as incorrect when the graded level of the rhythmicity or the non-rhythmicity was not predicted well enough (see Fig.4i,iii). For example, in rec.#11 (Fig.4i), the 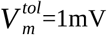 value did not display a high enough value to be linearly related to NSI_pLFP_ level according to the relationship *F*.

Using the tolerance criteria defined in the above paragraph, the accuracy of detection of a V_m_-based NSI value from the pLFP-based NSI value was 79.7±10.2% (mean ± s.e.m. over the n=14 recordings and all validated episodes). Decreasing the tolerance parameter values to *p^tol^*=1μV and 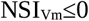 (corresponding to extremely strict matching criteria) led to an accuracy of 57.2±15.1%, meaning that, in more than half of the cases, the network state could be identified even with such remarkably high precision.

We noted that the misclassifications were not homogeneously distributed across different states (see Table 2). A specific set of network state misclassifications represented 65.3% of the misclassifications over the merged episodes across all recordings (i.e. 13.1% of all episodes). In this set, rhythmic activity predicted from the V_m_ (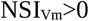) was misclassified as non-rhythmic activity from the pLFP (NSI_pLFP_>0). We analyze in the next section the reasons behind this finding.

**Table 1.** Parameters of the NSI_pLFP_ characterization. Note that the *p*_0_ and 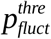 parameters are data-driven quantities, i.e. varying from recording to recording, deriving from the value of 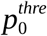 (reported as mean ± s.e.m over the n=14 recordings).

| Parameter | Symbol | Value |
| --- | --- | --- |
| pLFP root freq. | $f_0$ | 72.8 Hz |
| pLFP band factor | $w_0$ | 1.83 |
| pLFP smoothing | $T_{smoothing}$ | 42.2ms |
| percentile for pLFP lower bound | $p_0^{thre}$ | 1% |
| pLFP lower bound | $p_0$ | $2.85 \pm 0.73 \mu V$ |
| State window | $T_{state}$ | 400ms |
| Sliding mean window | $T_{mean}$ | 500ms |
| pLFP threshold for state validation | $p_{fluct}^{thre}$ | $2.85 \pm 0.73 \mu V$ |
| factor for rhythmicity threshold | $\alpha$ | 2.87 |

**Table 2.** Misclassifications in the pLFP-based (NSI_pLFP_) versus V_m_-based (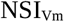) NSI characterization. Proportions of the misclassifications splitted into the rhythmic (NSI*≤*0) and non-rhythmic (NSI>0) cases for both the pLFP and V_m_ signals over all episodes in the n=14 recordings (related to Fig.4c).

| Misclassification cases | percentage |
| --- | --- |
| $NSI_{pLFP} > 0$ and $NSI_{Vm} \leq 0$ | 65.3% |
| $NSI_{pLFP} \leq 0$ and $NSI_{Vm} > 0$ | 25.0% |
| $NSI_{pLFP} > 0$ and $NSI_{Vm} > 0$ | 5.4% |
| $NSI_{pLFP} \leq 0$ and $NSI_{Vm} \leq 0$ | 4.3% |

### Graded aspect of network states in terms of pLFP and V_m_ fluctuations: rhythmic versus non-rhythmic states

The above analysis revealed a stronger tendency to classify network states as rhythmic from the V_m_ than from the pLFP fluctuations. This difference in classification suggested that the NSI measures revealed previously unexplored asymmetries between the characteristics of fluctuations of pLFP and V_m_ signal across different network states. We therefore analyzed more closely the correspondence between pLFP and V_m_ fluctuations for different levels of pLFP-based NSI and how this impacted our classification results

For non-rhythmic activity (NSI_pLFP_>0), the level of the NSI_pLFP_ was given by the mean pLFP deflection (Y-p_0_) in the time window T_state_=400ms. We thus compared the positive NSI values to the mean membrane potential depolarization level in the same window T_state_. For rhythmic activity (NSI_pLFP_≤0), the level of the network state index was proportional to the delta envelope of the pLFP. We thus compared the negative NSI values to the delta envelope in the V_m_ signal. We show the relationship between those pLFP-defined and V_m_-defined levels for two recordings in Fig.5a,b (for rec. #1 and rec. #11 respectively) and for the population data on Fig.5c.

**Figure 5.**
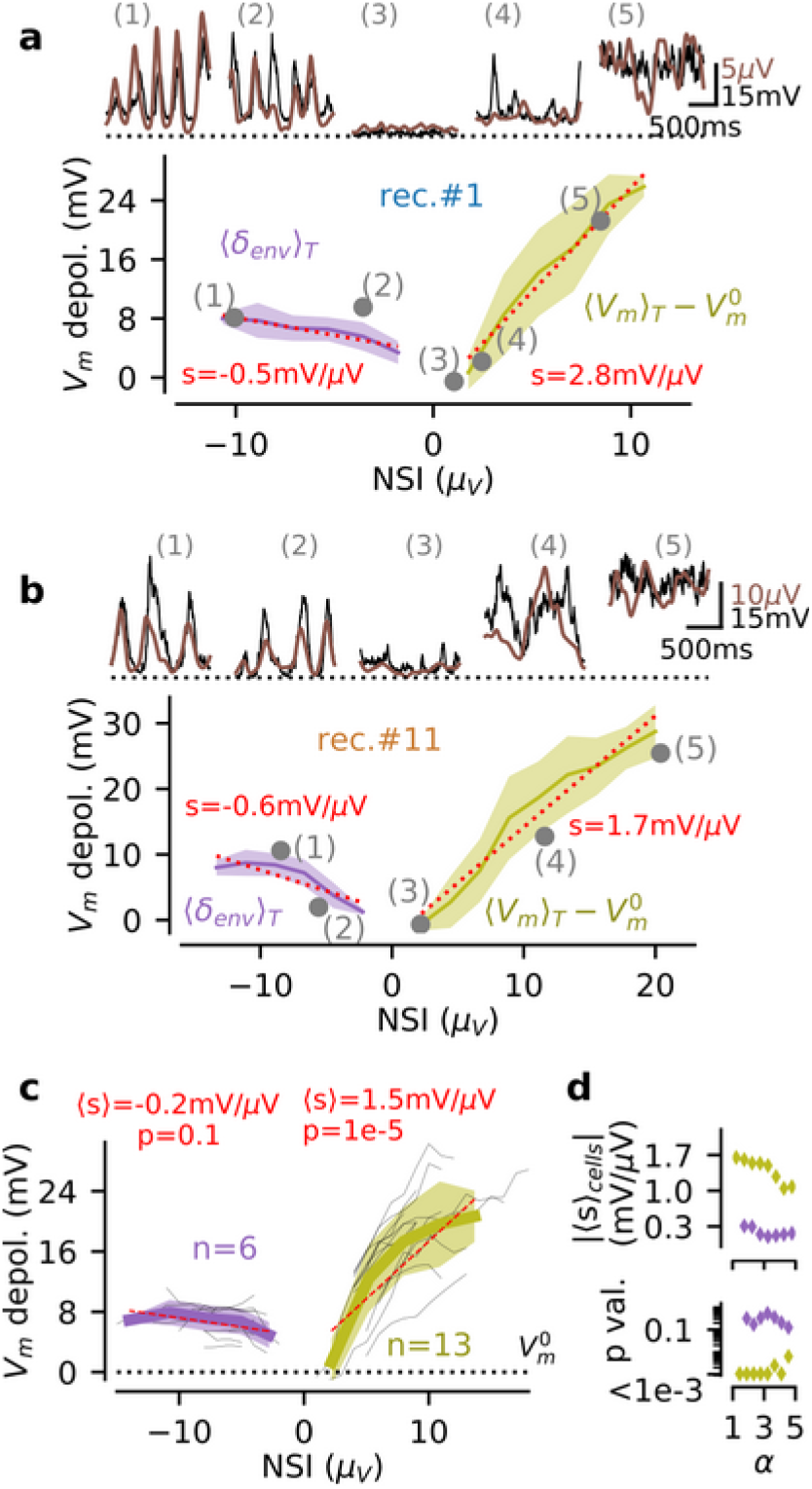
Relationship between pLFP-derived population activity (NSI_pLFP_) and single cell depolarization (V_m_) in rhythmic and non-rhythmic regimes. **(a)** Relationship, at the single recording level (shown for rec. #1), between the NSI_pLFP_ levels and the properties of the V_m_ fluctuations. For rhythmic states (NSI_pLFP_≤0, purple color), we show the relationship between the NSI level and the V_m_ amplitude of the delta-band envelope. For non-rhythmic states (NSI_pLFP_>0, kaki color), we show the link between the NSI level and the mean *V_m_* depolarization level over a *T_state_*=400ms window. We highlight with gray dots the values of the single episodes visible on Fig.3a (reproduced on the top inset, V_m_ in black and pLFP in brown). We show the linear regressions for the rhythmic and non-rhythmic data (dashed red line). Note that the plain curve does not reach episode 3 because the minimum number of episodes for averaging is not reached at that level (see Methods). **(b)** Same than **a** for rec.#11. **(c)** Reproducing the analysis of **a**,**b** over all n=14 recordings (see main text). We show the mean relations over recordings (thin gray lines) and the mean (wide curve) and standard deviation (shaded area) across recordings. Evaluated only for the NSI_pLFP_ levels displayed by multiple recordings (i.e. n=6 recordings for rhythmic activity and n=13 recordings for non-rhythmic activity). For each recording, we perform a linear regression with respect to the NSI_pLFP_ levels, we compute the mean across recordings 〈s〉 and the probability of a deviation from the 0-slope hypothesis *p* (paired t-test). **(d)** Relationship between the α parameter (that sets the proportion of non-rhythmic episodes, see **Fig. 3c**) and the mean slope over cells (top, 〈s〉 cells) together with the p-value testing the significance of a non-zero slope (bottom), i.e. reproducing the analysis of **c** for different α values.

Overall, we found (Fig. 5) that the membrane depolarization exhibited a strong dependency on the pLFP-based NSI level for non-rhythmic activity (NSI_pLFP_>0). We showed single episodes of increasing NSI_pLFP_ levels (numbered 3,4,5,6 in Fig.5a,b) and their respective episode averages at all NSI_pLFP_>0 levels for those two recordings (kaki curves in Fig.5a,b). The ~15μV variability in terms of pLFP based NSI levels corresponded to a ~30mV variability of depolarization level with a clear monotonic relationship for those two sample recordings. This behavior was confirmed at the population level (Fig.5c). We standardized the analysis across all recordings by binning the NSI_pLFP_ levels from 0μV to 30μV (a range covering all observed values) in bins of 1μV. We found that all recordings exhibited depolarizations with a steep dependency on the NSI_pLFP_ level 1.5±0.7mV/μV, significantly deviating from the null hypothesis of a zero slope (p=1.2e-5, n=13 recordings, paired t-test). On the other hand, we found that the value of pLFP-based NSI for rhythmic activity (NSI_pLFP_ ≤0) had a much lower impact on the membrane depolarization level, see Fig.5. The ~10μV variability in terms of pLFP based NSI levels translated into a weakly modulated V_m_ oscillation with a 5-10mV amplitude (purple curves in Fig.5a,b). We highlight this weak dependency on the selected samples shown in Fig.5a,b. We extended the analysis to all recordings, after standardizing the data by binning the pLFP-based NSI levels from −30μV to 0μV. At the population level, we confirmed that the mean membrane depolarization had a weak dependency on the pLFP-based NSI level (−0.2±0.3mV *μ*V), which was not significantly deviating from the null hypothesis of a zero slope (p=0.14, over the n=6 recordings displaying rhythmic activity within multiple NSI_pLFP_ bins, paired t-test). It should be noted that the lack of graded V_m_ variations for rhythmic activity was not related to our “rhythmicity threshold” limiting the set of rhythmic samples to a potentially-biased subset. When varying the rhythmicity-factor α up to α=5 (where the occurrence of rhythmic NSI_pLFP_ ≤0 states reaches ~80%, see Fig.3b), the depolarization level still showed a very weak dependency to the NSI level compared to that for non-rhythmic activity (Fig.5d, top) with similar statistical significance values (see Fig.5d, bottom).

We concluded that the graded levels of neural activity captured by the NSI_pLFP_ had a strong correlate in terms of membrane potential depolarization for non-rhythmic activity (NSI_pLFP_>0), while the various pLFP-based NSI levels of rhythmic activity (NSI_pLFP_≤0) rather corresponded to a stereotypical V_m_ oscillation with ~5-10mV amplitude.

Those observations explained the results of our cross-validation analysis (Table 2). The high precision of the classification in the jointly non-rhythmic case (“NSI_pLFP_>0 and 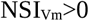” in Table 2, only 5.4% of the misclassifications) resulted from the strong relationship between the pLFP and V_m_ signals during non-rhythmic states (Fig.5a-c, kaki curves). On the other hand, the existence of a few episodes showing a high envelope delta oscillation in the V_m_ with a low delta envelope in the pLFP (cases such as episode 2 in Fig. 5a,b) created an ambiguous situation for the classifier because rhythmicity was hard to establish from the pLFP signal in those episodes. The cases with mixed rhythmic/non-rhythmic predictions indeed represented 90.3% of the misclassifications (“NSI_pLFP_ >0 and 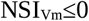” and “NSI_pLFP_ ≤0 and 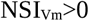” in Table 2). In particular, predicting rhythmicity from the V_m_ and non-rhythmicity from the pLFP was the prevalent misclassification case (65.3% for the case “NSI_pLFP_ ≤0 and 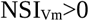”), consistent with the stronger representation of the delta pattern in the V_m_ than in the pLFP (NSI_pLFP_ ≤0 range in Fig. 5a-c). We confirmed that such misclassification cases originated from episodes of low delta envelope in the pLFP signal: the mean δ_env_ in misclassified cases was significantly lower than in accurately classified cases (1.5±0.3μV versus 2.7±0.5μV, p=3.3e-10 paired t-test, in the n=13 recordings displaying both conditions).

### Using the NSI_pLFP_ to quantify network state distributions in the somato-sensory cortex of awake head-fixed mice

We analysed the NSI_pLFP_ state distribution in the whole dataset (n=14 simultaneous V_m_ and LFP recordings in the somato-sensory cortex of awake head-fixed awake mice). Fig. 4 shows eight example recordings representative of the dataset, spanning the full range of correlations between the pLFP and V_m_ fluctuations (see Fig.2c). We show a 60s sample of the intra- and extracellular recordings together with the time-varying NSI_pLFP_ (Fig.4a) and the histogram of the NSI_pLFP_ levels across the whole recording (Fig.4b).

Overall, the dataset was dominated by non-rhythmic activity with a fraction of non-rhythmic states: *F*_*NSI* >0_ =81.1±15.9%. The fraction of rhythmic activity exhibited a large variability with n=3 recordings below *F*_*NSI* ≤ 0_=2% and n=3 recordings above *F*_*NSI* ≤ 0_=35% (peaking at *F*_*NSI* ≤ 0_=46.8% for recording #1, see Fig.4b).

The mean absolute NSI_pLFP_ value over rhythmic periods (i.e. NSI_pLFP_≤0) was 5.65±1.02μV (mean ± s.e.m over n=14 recordings) and 6.67*±*1.34*μ*V for non-rhythmic periods (i.e. NSI_pLFP_>0), yielding a significant increase from rhythmic to non-rhythmic states (p=3.0e-5, paired t-test). The increased amplitude in terms of pLFP signal (i.e. high frequency content of the LFP) suggested that, as an average, synaptic activity was stronger in the non-rhythmic periods that in the rhythmic ones (see Discussion). This increased range of pLFP level was also true for the maximum level displayed by single recordings with 9.52±2.25μV versus 12.06±3.4μV (p=9.8e-3, paired t-test) for rhythmic and non-rhythmic activity, respectively. During non-rhythmic periods, we found a mild but significant increase of the variability (standard deviation, from 1.29±0.32μV to 1.56±0.71μV, p=8.9e-2, paired t-test) and skewness of the pLFP distributions (from 0.35±0.33 to 0.7±0.41, p=8.2e-2, paired t-test).

### Network state variability within individual recordings predicts the average correlation between population and single cell signals

We next analysed whether the NSI_pLFP_ state distributions could explain the variability of the correlation cc(V_m_, pLFP) between the time-varying population signal pLFP(t) and the single cell signal V_m_(t) in our recordings (see Fig. 4a,b). We reduced the NSI_pLFP_ distribution *per* recording to a few components (detailed below) and we used univariate and multivariate linear regressions to analyse how the variability of those components across recordings explained the variability of the observed correlation values.

From top to bottom in Fig.4b (i.e. from high to low correlation recordings), the distribution of NSI_pLFP_ levels across recordings was quantitatively different. At the top (high correlations, cc(V_m_, pLFP)>0.4, panels i-v), distributions were bimodal with two high peaks both in the rhythmic (NSI_pLFP_≤0) and non-rhythmic (NSI_pLFP_>0) areas. At the bottom (low correlations, cc(V_m_, pLFP)<0.4, panels vi,vii), distributions were dominated by non-rhythmic episodes with rather narrow range of NSI_pLFP_ values within the (NSI_pLFP_>0) domain. To quantify these features within the single recording distributions, we decomposed each distribution into the following quantities: *i)* the mean NSI_pLFP_>0 over the whole recording μNSI; *ii)* the variability (standard deviation) of the full NSI distribution σ_NSI_; *iii)* the fraction of rhythmic episodes *F*_*NSI* ≤ 0_; *iv)* the mean in NSI_pLFP_ values restricted to non-rhythmic episodes *μ*_*NSI* > 0_; *v)* the mean in NSI_pLFP_ values restricted to rhythmic episodes *μ*_*NSI* ≤ 0_; *vi)* the variability in NSI_pLFP_ values restricted to non-rhythmic episodes *σ*_*NSI* > 0_ ; vii) the variability in NSI_pLFP_ restricted to rhythmic episodes *σ*_*NSI* ≤ 0_.

We show the results of a linear regression analysis for each factor in Fig.6a (ordered by explained variance) and their relationship with the correlation value in Fig.6b. The factor explaining the highest percentage of the variability (55.27% of the full variability) was the standard deviation of the NSI distribution *σ_NSI_*. Strikingly, the fraction of (synchronous) rhythmic states was only the third most important factor with an explained variance of 33.89%. A more important factor was found to be the variability of network states within non-rhythmic states *σ*_*NSI* > 0_ with an explained variance of 41.5%. Recording #14 in Fig.4ii provided an example of these observations. It exhibited strong variability in terms of non-rhythmic states and relatively low occurrence of rhythmic activity (e.g. lower than recording #1). However, it still displayed high correlation coefficient (with cc(*V_m_*, pLFP)=0.66). Other individual factors had weak statistical significance (p>0.04, see Fig.6a). However, using a multiple linear regression including all factors, we found that different components of the NSI distributions had complementary contributions in shaping the average correlation *per* recording. The full linear model indeed yielded an explained variance of 69.17% (after correction by the number of linear factors, f-test p=3.1e-2). Reduce the dimensionality of the linear model (up to three components), we found that combining the significant sub-components of the NSI variability (*σ*_*NSI* > 0_ and *F*_*NSI* ≤ 0_) with the variability within rhythmic states *σ*_*NSI* ≤ 0_ produced a statistically-significant model (f-test, p=6.6e-3) predicting 59.84% of the variability (corrected by the number of factors).

**Figure 6.**
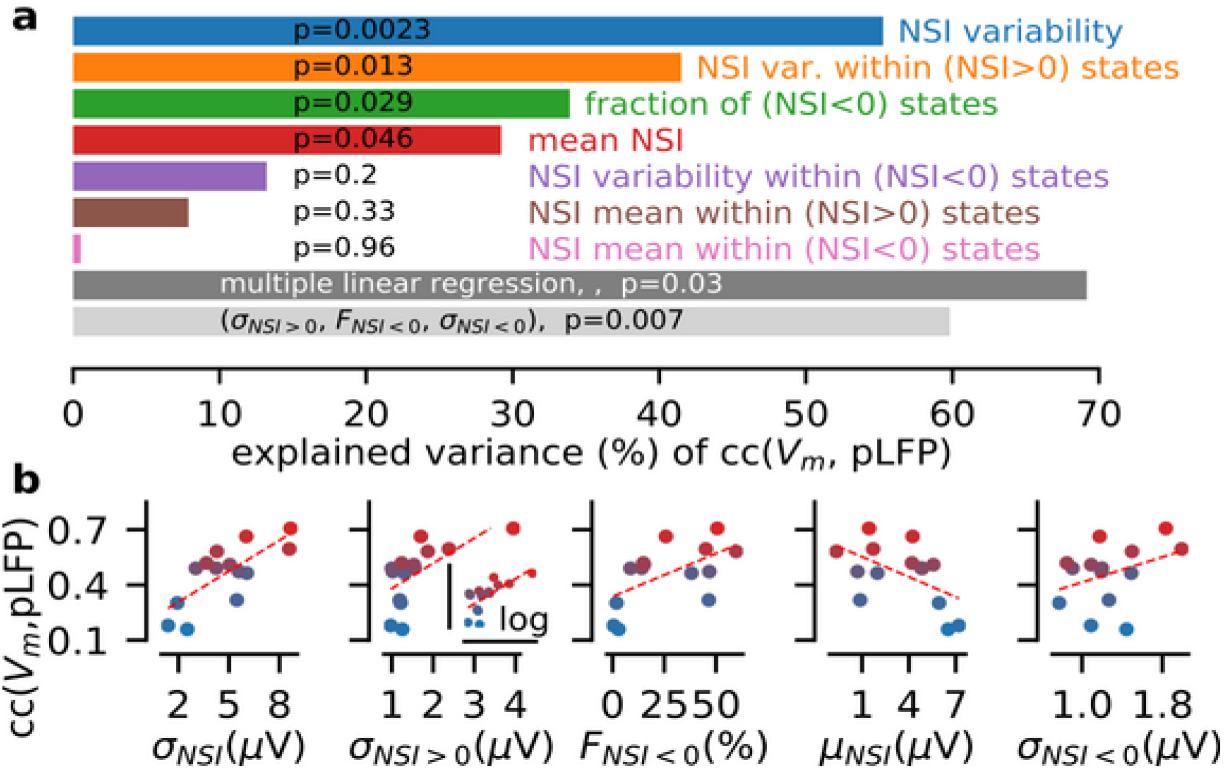
Features in the distribution of the pLFP-based NSI explain the diversity over recordings of the correlation value between the population (pLFP) and single cell (V_m_) signals. **(a)** We perform a linear-regression between the correlation coefficient cc(V_m_,pLFP) and the following quantities characterizing the NSI_pLFP_ distribution: 1) the variability σ_NSI_ (standard deviation) of the NSI_pLFP_ values across the whole recording, 2) the variability in NSI_pLFP_ values restricted to non-rhythmic episodes σ_NSI>0_, 3) the fraction of rhythmic episodes F_NSI≤0_, 4) the mean NSI_pLFP_ over the whole recording μ_NSI_, 5) the variability in NSI_pLFP_ values restricted to rhythmic episodes σ_NSI≤0_, 6) the mean in NSI_pLFP_ values restricted to non-rhythmic episodes μ_NSI>0_, 7) the mean in NSI values restricted to rhythmic episodes μ_NSI≤0_. We show the explained variance for all quantities on the x-axis and the statistical significance of those linear models (p-values). We also performed multiple linear regressions with all those factors (dark gray bar) and the best three-component model (light gray bar: σ_NSI>0_, F_NSI≤0_, σ_NSI≤0_). We report the variance corrected by the number of linear factors (adjusted *R^2^*). **(b)** Scatter plot between the value of a given factor and the V_m_-pLFP correlation coefficient across individual recordings. Shown for the first five factors of the NSI_pLFP_ distribution with the highest percentage of explained variance (see variance explained and p-values in **a**). The color code of individual recordings matches that of Fig.2c.

We concluded that, in the present dataset, the average correlation between a single cell signal (V_m_) and the population signal (pLFP) could be largely explained (up to ~70%) by features of the NSI_pLFP_ distributions. State variability among all NSI_pLFP_-defined states (*σ_NSI_*) was a critical factor in determining such a variability (blue in Fig. 6a). We decomposed this state variability and found that the two most prominent factors are the variability within non-rhythmic states (*σ*_M*NSI* > 0_ , orange in Fig 6a) followed by the occurrence of synchronous rhythmic activity (*F*_*NSI* ≤ 0_, green in Fig. 6a).

## Discussion

In this study, we developed a method to extract from LFP recordings in the awake mouse cortex network states information that previously could be obtained only with intracellular V_m_ recordings (Arroyo et al., 2018; Einstein et al., 2017; McGinley et al., 2015a; Nestvogel and McCormick, 2021; Polack et al., 2013; Poulet and Petersen, 2008; Reimer et al., 2014). Prolonged membrane potential recordings are difficult to achieve, and most of the times require head-fixation (but see Lee et al., 2006). This strongly limits our ability to describe the complexity of network states and their relationship with behavior. Achieving precise state classification based on LFP recordings, which are technically easier to perform and can be performed in freely moving animals, will greatly increase our ability to understand the cellular and network mechanisms underlying cortical processing during behavior. Previous attempts to classify network states from the LFP have been limited to the delta-band activity (Chen et al., 2017; Pala and Petersen, 2018; Vinck et al., 2015). When applied to data gathered in awake animals, the delta-to-gamma state classification used in anesthetized preparations (Cheng-yu et al., 2009; Saleem et al., 2010) was shown to only separate between the two extremes of the spectrum of cortical states: synchronized delta-band activity and desynchronized activity at high gamma power. In contrast, the new classification method developed in this study, NSI, captured the large spectrum of network states in the awake neocortex. Importantly, it provided quantitative measurement of the “U-model” of cortical states, which was previously developed only based on intracellular membrane potential recordings (McGinley et al., 2015b).

Prior to the NSI classification, we found it essential to apply a pre-processing step to the extracellular LFP. We computed the time-varying envelope after a wavelet transformation in the high-gamma band (yielding the pLFP signal). In a previous study in cat neocortex (Mukovski et al., 2006), the authors identified active and silent states (Up and Down respectively) under anaesthesia from the pLFP signal (with a slightly different frequency band of the LFP: 20-100 Hz, instead of the data-driven [39.7,133.6] Hz band used here). We showed that such a signal processing step enabled capturing various states of wakefulness, ranging from delta (~3 Hz) oscillatory activity to desynchronized states at various levels of spiking activity.

We then used the NSI to analyze how the distribution of network states within a recording period shaped the average correlations between the single cell (intracellular) and population (extracellular) signals. In anesthetized preparations, a consolidated view suggests that low frequency activity is the main source for neural synchrony in cortical networks and thus for cell-to-population correlation (Steriade et al., 1993). Also in our dataset recorded during wakefulness, the fraction of slow (delta-band) oscillatory episodes was a factor significantly contributing to the level of correlation between single cell and population signal (Fig. 6). However, and unexpectedly, we found that this was a weaker factor than the variability in the set of non-rhythmic states NSI values of cortical activity (Fig. 6). This observation was explained by the fact that synchronized delta activity represented only a modest fraction of network activity during wakefulness (18.9% here) and by the fact that the diverse non-rhythmic states corresponded to strongly differing levels of both the V_m_ depolarization and the high-gamma activity (Reimer et al., 2014; McGinley et al., 2015a; Zerlaut et al., 2019, Fig. 2,3,4). Importantly, the diversity of network state distributions across recordings largely contributed to the high variability in the measured cell-to-population correlations (70% of this variability could be explained by recording-specific features of the NSI_pLFP_ distribution, Fig. 6). This observation further highlights the necessity of network-state monitoring in the interpretation of experimental results in the awake cortex (McGinley et al., 2015b), and it suggests the possible importance of NSI indices to understand and characterise how the degree of coupling between single-cell and population-level activity varies across different behavioural states.

Our results provide possible insights on the circuit dynamics during different cortical states observed during wakefulness. First, non-rhythmic episodes had population activity levels (pLFP) varying over a wide range and seemed to reach up to levels never observed in rhythmic episodes (Fig 3f). Second, in such non-rhythmic states, we observed a tight relationship between the population activity and single-cell depolarization (Fig. 5). This suggests that local recurrent spiking activity plays a major role is shaping the dynamics of non-rhythmic states. It also corroborates, at the level of population signals, that the hyperpolarized non-rhythmic states observed at intermediate arousal when sensory detection is optimal are characterized by lower levels of ongoing recurrent synaptic activity in local cortical populations (McGinley et al., 2015; Neske et al., 2019; Nestvogel and McCormick, 2021). Instead, during rhythmic states, population activity varied over a much more limited range, and the relationship between population activity (pLFP) and single-cell membrane potential was much less tight (Fig. 5). This suggests that in rhythmic states local recurrent cortical activity plays a lesser role in shaping cortical dynamics when compared to non-rhythmic states. This view is consistent with the critical role of the thalamus in shaping ~3Hz oscillatory activity in the cortex (Nestvogel and McCormick, 2021). Because of the occasionally low level of the pLFP signal during rhythmic activity, the NSI_pLFP_ was biased towards low activity non-rhythmic states when evaluated from extracellular recordings. Despite this limitation, the overall high matching value obtained through cross-validation (~80% correct) suggested that the pLFP-based NSI is a valid index to characterize the different network states occurring during wakefulness.

Given the widespread use of extracellular recordings in neuroscientific research in both head fixed and freely moving preparations (Jun et al., 2017; Panzeri et al., 2015) and the relevance of state modulation in sensory processing (Arandia-Romero et al., 2016; Ayaz et al., 2013; Busse et al., 2017; Bennett et al., 2013; Dadarlat and Stryker, 2017; Davies et al., 2020; Destexhe, 2011; Ecker et al., 2014; Einstein et al., 2017; Fu et al., 2014; Lee et al., 2014; ; Muller et al., 2018; Niell and Stryker, 2010; Pachitariu et al., 2015; Pakan et al., 2016; Pinto et al., 2013; Polack et al., 2013; Poulet and Crochet, 2018; Reimer et al., 2014; Vijayan et al., 2010; Vinck et al., 2015; Zhou et al., 2014), the presented method provides a potentially important analytical tool to document the properties and functions of state-dependent computations in neocortex (Buonomano and Maass, 2009; Cardin, 2019).

## Material and Methods

### Animals

Experimental procedures involving animals have been approved by the IIT Animal Welfare Body and by the Italian Ministry of Health (authorization # 34/2015-PR and 125/2012-B), in accordance with the National legislation (D.Lgs. 26/2014) and the European legislation (European Directive 2010/63/EU). Experiments were performed on young-adult (4-6 weeks old, either sex) C57BL/6J mice (Charles River, Calco, Italy). The animals were housed in a 12:12 hr light-dark cycle in singularly ventilated cages, with access to food and water ad libitum.

### Experimental design

The experimental procedure for simultaneous extra- and intra-cellular recordings in awake head-fixed mice have been previously described (Zucca et al., 2017) and the present dataset was used in a previous study (Zerlaut et al., 2019). Briefly, a custom metal plate was fixed on the skull of n=4 young (P22-P24) mice two weeks before the experimental sessions. After a 2-3 days recovery period, mice were habituated to sit quietly on the experimental setup for at least 7-10 days (one session per day and gradually increasing session duration). The day of the experiment, mice were anesthetized with 2.5% isoflurane and a small craniotomy (0.5×0.5 mm) was opened over the somatosensory cortex. A 30-minute long recovery period was provided to the animal before starting recordings. Brain surface was kept moist with a HEPES-buffered artificial cerebrospinal fluid (aCSF). Local field potential (LFP) recordings were performed by lowering a glass pipette filled with aCSF into the tissue with the tip placed at ~300 μm from pial surface. Simultaneous current-clamp patch-clamp recordings were carried out on superficial layers (100– 350 μm), all recorded cells had a regular-spiking response to current pulses (data not shown) and were therefore identified as putative pyramidal neurons (Connors and Gutnick, 1990). 3–6 MΩ borosilicate glass pipettes (Hilgenberg, Malsfeld, Germany) were filled with an internal solution containing (in mM): K-gluconate 140, MgCl2 1, NaCl 8, Na2ATP 2, Na3GTP 0.5, HEPES 10, Tris-phosphocreatine 10 to pH 7.2 with KOH. Current-clamp recordings were not corrected for liquid junction potential offset. Electrical signals were grounded at the top of the skull (at the location of the craniotomy) and were acquired using a Multiclamp 700B amplifier, filtered at 10 kHz, digitized at 50 kHz with a Digidata 1440 and stored with pClamp 10 (Axon Instruments, Union City, CA). Multi-Unit Activity (MUA) was computed by band-pass filtering (0.3-3 kHz) the extracellular signal and taking the absolute value of the resulting signal (Einevoll et al., 2013).

### Wavelet Transform

Our signal processing pipeline of the LFP was based on the wavelet transform. We implemented a wavelet transform based on the Morlet wavelet, which has the following equation (Mallat, 1999):

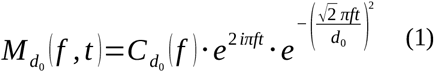

where *f* is the frequency of the wavelet and *d*_0_ the decay parameter of the envelope. We used a value of *d*_0_=6 throughout the study. The coefficient 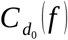 is the normalization coefficient of the wavelet. Note that, to keep a meaningful link with the physical units of the signal, we did not normalize the wavelet with respect to itself, but with respect to a sinusoid (otherwise the wavelet transform with standard normalization of a sinusoid of frequency *f* and amplitude 1 has a value greater than 1 at the frequency *f*). The wavelet normalization coefficient was therefore defined, for a wavelet frequency *f* and an extent *Ω*, as:

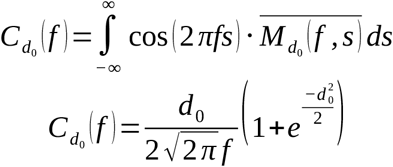

The transform was implemented by a convolution between the complex conjugate of the Morlet wavelet (Eq.1) and the signal *S* (*t*), i.e.:

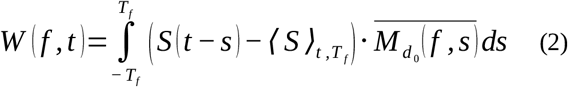

where 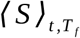 is the signal average in the window centered at *t* of extent *T_f_*. *T_f_* is the frequency-dependent window on which the convolution is performed, it was defined as the extent of the wavelet where its amplitude decays by (1 − *e*^−4^)=98.2%, i.e. 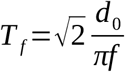. From Eq.2, we computed the envelope and the phase at time *t* of a given frequency *f* in the signal by taking the norm and argument of the complex number *W* (*f, t*).

### Computing the pLFP signal

From the LFP time series, we computed a “processed LFP” (Mukovski et al., 2006), shortened to “pLFP”, which corresponds to the temporally-smoothed high-gamma envelope variations of the LFP fluctuations (see main text). The pLFP was computed as follows. We consider a frequency band spanning [*f*_0_ / *w*_0_, *f*_0_⋅ *w*_0_], where *f*_0_ is a root frequency and *w*_0_ the width parameter of the band. We take a set of *N* =5 wavelets uniformly spanning this band (i.e. evenly space from *f*_0_ / *w*_0_ to *f*_0_⋅ *w*_0_). The pLFP signal was computed as the sum over *N* of the *k* wavelet envelopes of frequency *f_k_* ∈ [*f*_0_ / *w*_0_ , *f*_0_⋅ *w*_0_], i.e.:

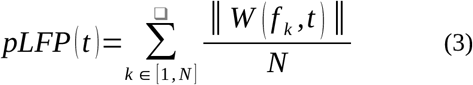

This time-varying signal is then smoothed over time with a Gaussian filter to yield the final pLFP signal (see Fig.2a). The parameters *f*_0_, *w*_0_, and the time smoothing width *T_smooth_* were set to maximise the correlations between the time course of the membrane potential and the pLFP (see Results) and their value is reported in **Table 1**.

### Computing the Network State Index (NSI)

Then, from the pLFP we computed the Network State Index (NSI), as follows. The pLFP was first downsampled by averaging over bins of 1ms (keeping the 50kHz sampling of electrophysiological signals is unnecessary given the much slower time scale of state transitions, see Results). We then computed the distribution of the pLFP over the whole recording and we extract the baseline noise level of the pLFP signal *p*_0_ by taking the value of the first (lowest) percentile of the distribution. We interpret this recording-specific *p*_0_ value as the residual high-gamma envelope in absence of neural activity (see the systematic depolarization from the activity at pLFP≤p_0_ in Fig.3f) and we therefore considered it as an estimate of the noise level in the pLFP signal.

We computed the time-varying envelope of the [2,4] Hz band *δ_env_* of the pLFP signal using the above wavelet transform (Eq.2). We took a set of 20 wavelets uniformly sampling the [2,4] Hz band and, at every 1ms time point, we extract the maximum envelope from this band (and the phase of this maximum envelope in Fig.3f). We constructed a weighted estimate *X* (*t*) of the low-frequency content of the pLFP signal by *X*(*t*) = *p*_0_+*α δ_env_*, where *α* is the threshold parameter for the rhythmic/non-rhythmic classification (see Results). We also computed a slow average of the pLFP fluctuations with a Gaussian smoothing of time constant *T mean*=500ms, yielding the signal *Y* (*t*).

Finally, the Network State Index (NSI) was defined from the above computed signals by the following equation:

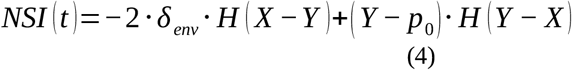

where *H* is the Heaviside step function.

From this time-varying signal, we computed what we termed “validated” network states by running through the time axis in steps of *T_state_*/2=200ms and identifying those time periods in which t the *T_state_*=400ms window surrounding each time point does not contain variations of the NSI signal larger than the noise level *p*_0_. When averaging quantities for a given NSI level (Figure 3,4,5), we considered only the NSI levels including more than 5 validated episodes to get a meaningful average.

The signal processing steps of this procedure are illustrated on Fig.3a and all parameters of the analysis are summarized in Table1.

### Statistical analysis

Experimental data were imported in Python using the neo module (Garcia et al., 2014). All signal processing steps (sub-sampling, convolution, filtering) were implemented in numpy (Harris et al., 2020). Statistical analysis was performed with the *scipy.stats* module of SciPy (Oliphant, 2007). We analysed the linear relationship between continuous samples with a Pearson correlation analysis (function *scipy.stats.pearsonr*) and we reported the two-tailed p-value of the null correlation hypothesis. For the statistics of samples consisting of an averages over a given recording session (n=14 recordings sessions), we tested the significance using two-tailed t-tests (functions *scipy.stats.ttest_rel*, *scipy.stats.ttest_ind* or *scipy.stats.ttest_1samp* for paired samples, unpaired samples and single samples respectively). The multiple linear regressions of Fig. 6 was performed with the OLS (ordinary least squares) function of the *statsmodel* module. We analysed the statistical significance of the single or multi-component models with an F-test and we report the variance adjusted by the numbers of factors.

### Software Accessibility

We implemented the described analysis into a software publicly available at the following link https://github.com/yzerlaut/Network_State_Index. The software contains the described algorithm, a graphical user interface and the support for a few electrophysiological data formats.

## Acknowledgments

This research was supported by the Flag-Era JTC Human Brain Project (SLOW-DYN), by NIH BRAIN Initiative grants R01NS109961, NS108410, and U19NS107464, by the ERC (NEURO-PATTERNS 647725), and by the Fondation pour la Recherche Médicale (ARF201909009117).

